## Supplementary figures for "Dynamical Latent State Computation in the Posterior Parietal Cortex"

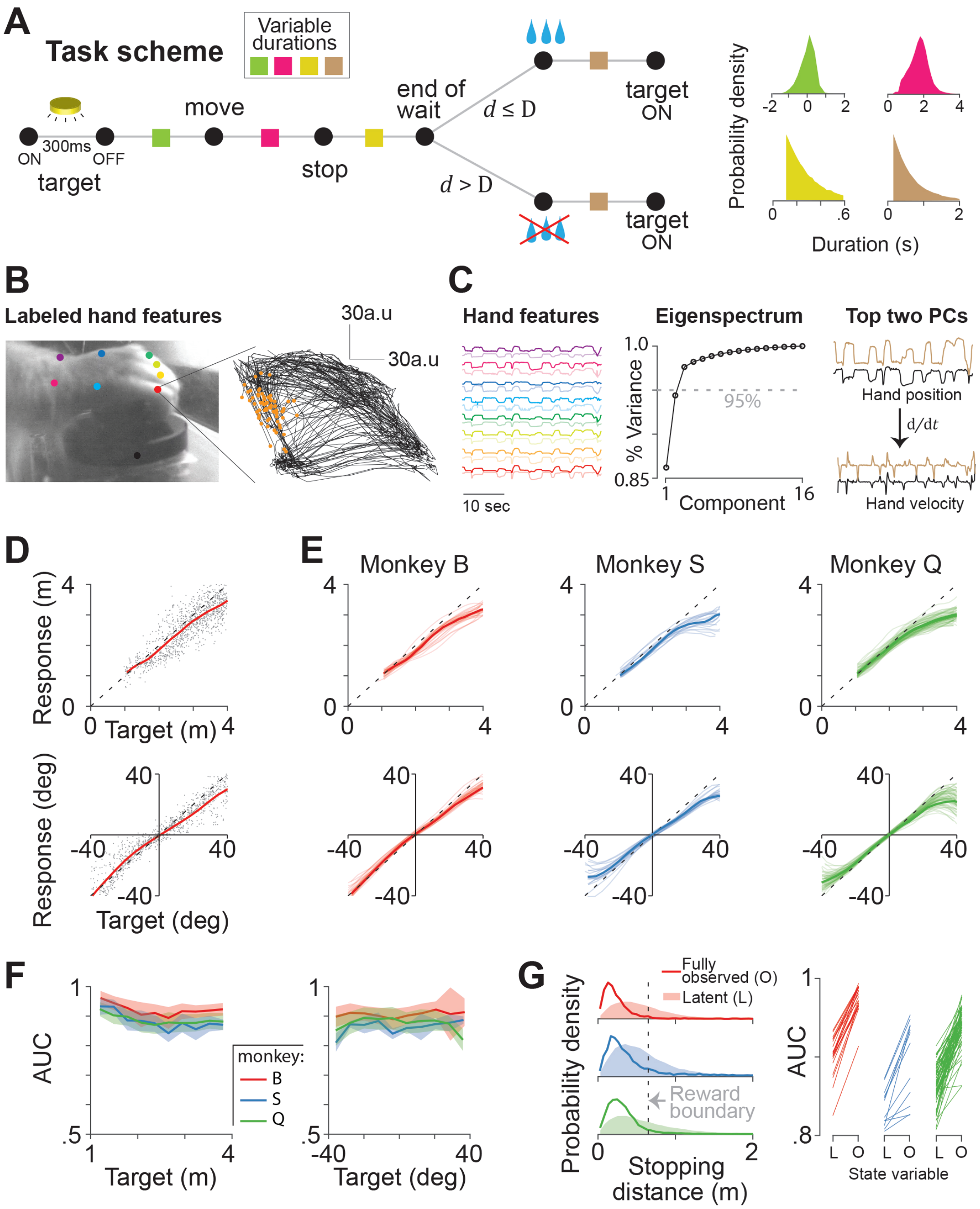
A
Task scheme
Variable durations
target
ON
300ms
OFF
move
stop
end of wait
d ≤ D
d > D
target ON
target ON
Probability density
Duration (s)
B
Labeled hand features
30a.u
30a.u
C
Hand features
10 sec
Eigenspectrum
% Variance
95%
Component
Top two PCs
Hand position
d/dt
Hand velocity
D
Response (m)
Target (m)
Response (deg)
Target (deg)
E
Monkey B
Monkey S
Monkey Q
F
AUC
Target (m)
Target (deg)
monkey:
B
S
Q
G
Probability density
Fully observed (O)
Latent (L)
Reward boundary
Stopping distance (m)
AUC
State variable

*Figure S1:* **A.** Left: The timeline of a typical trial: black circles denote salient events; colored squared denote the duration of intervening periods. Monkeys received juice rewarded if their position fell within the boundary of the reward zone at the end of the waiting period ($d$: distance from the target, D: radius of the reward zone). No juice was provided otherwise. Right: Empirical probability distributions of the duration of intervening periods shown in the left panel. Waiting period (dusty gold) and inter-trial interval (brown) were truncated exponential distributions with mean of 250ms and 500ms respectively. Movement onset times (green) and travel durations (hot pink) were entirely under the monkeys' control. Note that in order to allow behavior to be as natural as possible, monkeys were not forced to wait until the target disappeared, hence the negative durations in nearly $\sim$30% of trials (green). Monkeys were teleported to the starting position at the time of target onset, so movements initiated prior to target onset were inconsequential. **B.** Left: A random frame taken from the video recording of a monkey's hand movements while performing the task. Colored dots denote (eight) points that were labeled offline for the purpose of extracting hand coordinates using DeepLabCut. The position of the base of the joystick (labeled in black) served as a reference point for the coordinates. Right: A blow-up of the extracted spatial trajectory of one of the points (the little finger). **C.** Left: Spatiotemporal profile of hand movements: the time-course of the features extracted using the labeled points (8 points $\times$ 2 coordinates = 16 features) during a random 30 second segment of the video. Middle: The cumulative fraction of variance explained by the principal components (PCs) of the features (arranged in decreasing order of variance). Two leading PCs explain almost 95% of the total variance in the position of the hand. Right: Time-course of the top two PCs during the same epoch. The magnitudes of the time derivative of the components (i.e., hand speed in the principal subspace) were taken to be the motor output signal. **D.** Top: Comparison of the radial distance of the monkey's response (stopping location) against radial distance of the target across trials from one example session. Bottom: Angular eccentricity of the response versus target angle. Black dashed lines have unity slope. Red lines denote the locally linear regression (LLR) model fit to the data points. **E.** Thin lines show the LLR model fit to radial (top) and angular (bottom) responses from individual experimental sessions of all three monkeys. Thick lines denote the medians across sessions. **F.** Left: Area under the ROC curve (AUC) computed by binning trials based on the radial distance of the target. Right: Similar to left panel, but binning based on the angular eccentricity of the target. Shaded regions denote standard deviation across experimental sessions. **G.** Left: Probability distribution of stopping distances (from the target) for the set of trials in which the world state was either latent (shaded) or fully observable (open). Right: AUCs calculated based on the trials corresponding to the two conditions separately for individual sessions from all three monkeys (L – latent, O – fully observable).

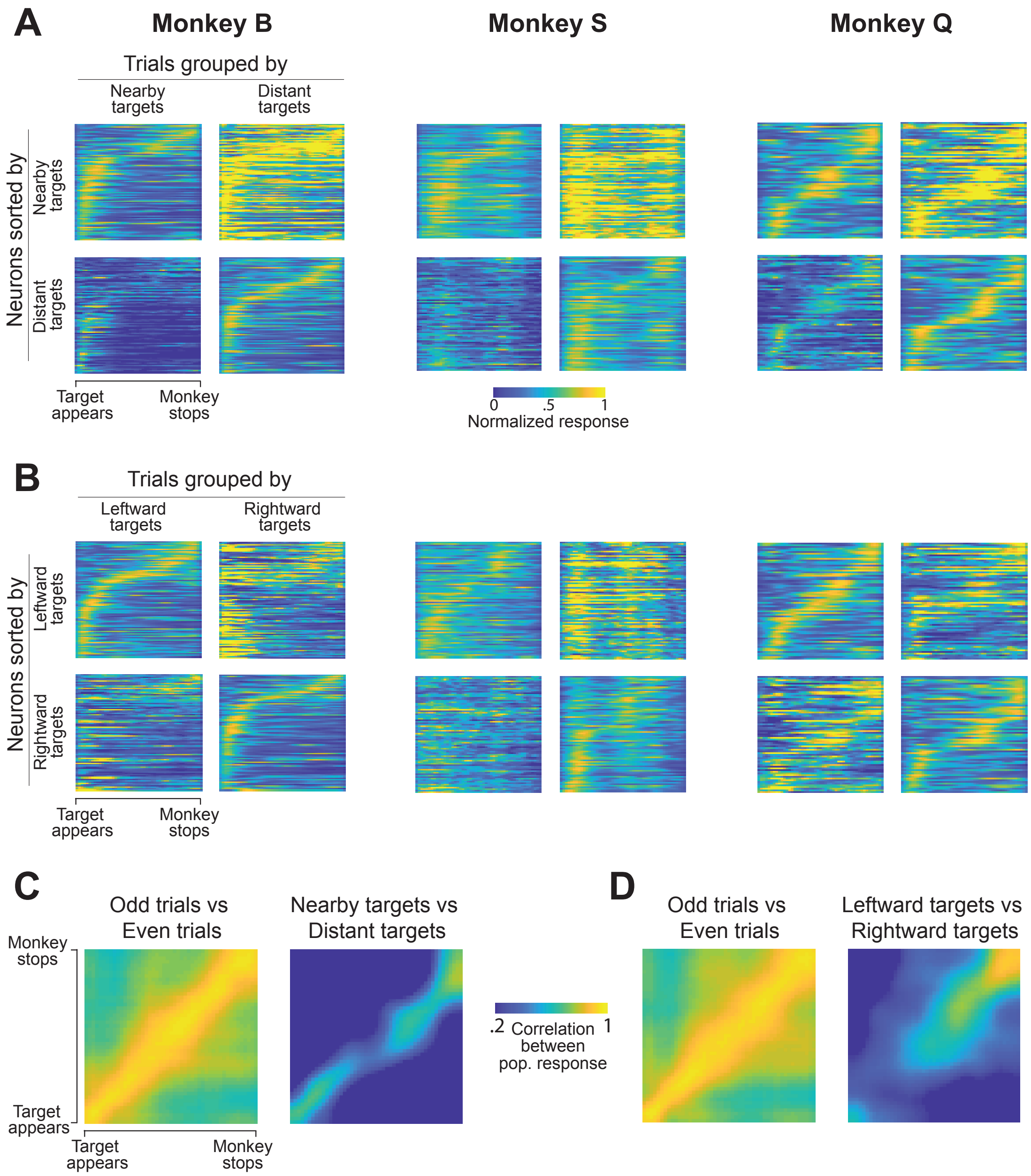
A
Monkey B
Monkey S
Monkey Q
Trials grouped by
Nearby targets
Distant targets
Neurons sorted by
Nearby targets
Distant targets
Target appears
Monkey stops
0
.5
1
Normalized response
B
Trials grouped by
Leftward targets
Rightward targets
Neurons sorted by
Leftward targets
Rightward targets
Target appears
Monkey stops
C
Odd trials vs Even trials
Nearby targets vs Distant targets
Monkey stops
Target appears
Target appears
Monkey stops
.2
Correlation between pop. response
1
D
Odd trials vs Even trials
Leftward targets vs Rightward targets

*Figure S2:* **A.** Peak-normalized response of neurons calculated by averaging across the set of trials with nearby (left panels) or distant (right panels) targets, separately from neural populations recorded simultaneously in three different monkeys. Neurons are sorted according to the timing of their peak response observed in the set of trials with nearby (top panels) or distant (bottom panels) targets. Spike times were rescaled based on the trial duration before trial-averaging and the resulting response profile of each neuron was subsequently normalized by the peak activity observed in the condition used for sorting. **B.** Similar to panel A, but with trials grouped according to the target angle. **C.** Correlation between population responses taken from trials within the same (odd vs even trials) or different (nearby vs distant targets) groups, computed for all pairs of time points. Note that the diagonal of this correlation matrix is the time course of pattern similarity shown in Figure 2C (right panel). **D.** Similar to left panel, but with trials grouped according to the target angle.

---

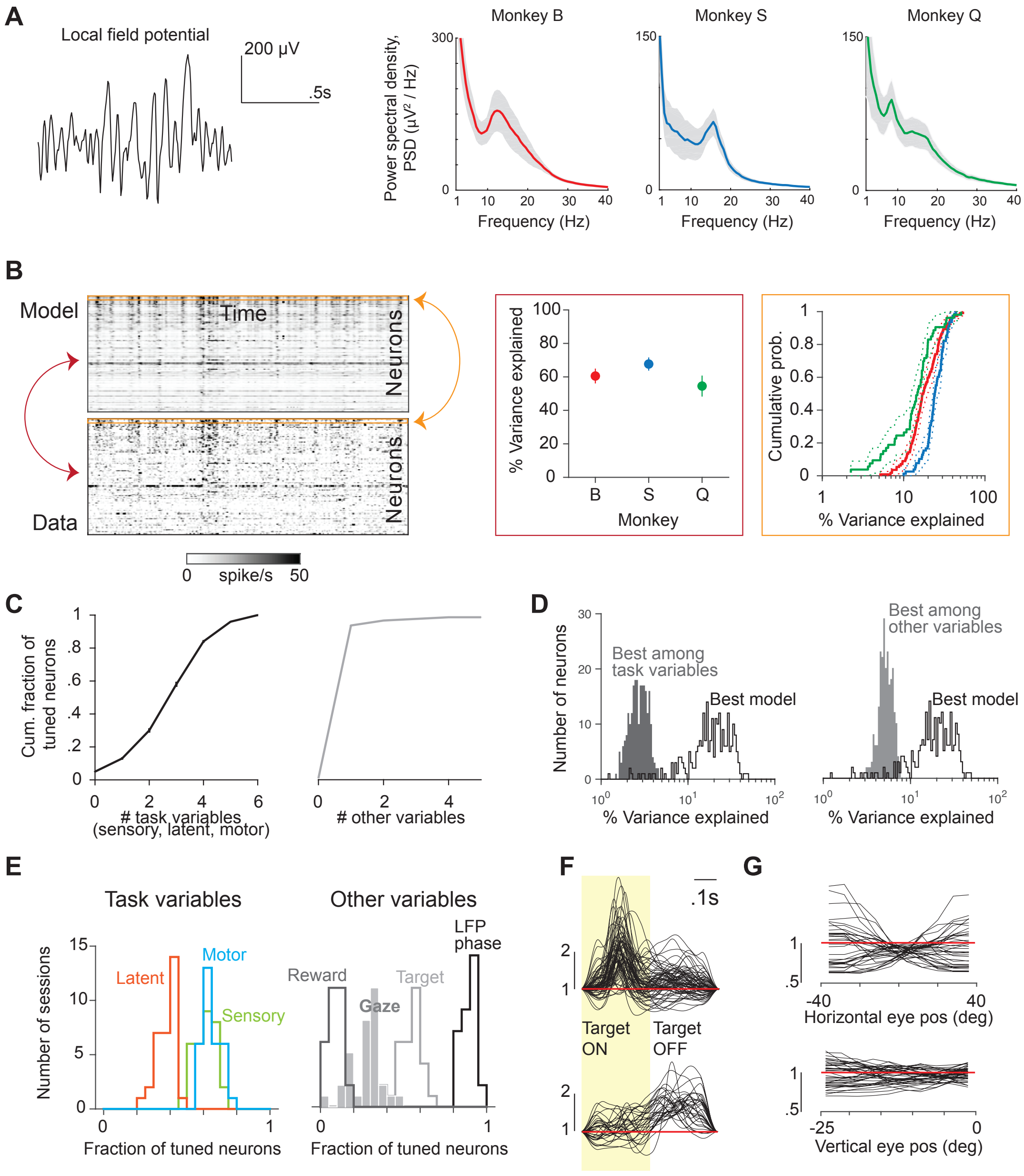
A
Local field potential
200 μV
.5s
Power spectral density, PSD (μV² / Hz)
Monkey B
Monkey S
Monkey Q
Frequency (Hz)
B
Model
Data
Time
Neurons
spike/s
% Variance explained
Monkey
Cumulative prob.
C
Cum. fraction of tuned neurons
### task variables (sensory, latent, motor)
### other variables
D
Number of neurons
Best among task variables
Best model
Best among other variables
E
Task variables
Other variables
Number of sessions
Latent
Motor
Sensory
Reward
Gaze
Target
LFP phase
Fraction of tuned neurons
F
.1s
Target ON
Target OFF
G
Horizontal eye pos (deg)
Vertical eye pos (deg)

*Figure S3:* **A.** Snippet of raw local field potential (LFP) (left) and power spectral density of LFPs recorded from different monkeys (right). **B.** Reproduction of 3C (left): Activity of simultaneously recorded neurons during a random thirty second epoch during the experiment (bottom) and the corresponding prediction reconstructed using the model (top). Neurons are arranged according to their contribution to the leading principal component (bottom – lowest; top – highest) for visualization. Percent of variance in the population activity structure (middle, maroon box) and single neuron activity (right, orange box) explained by the best-fit generalized additive model for data from each monkey. **C.** Left: Cumulative fraction of neurons as a function of the number of significantly predictive task variables (left) and significantly predictive other variables (right). Other variables include LFP phase, target onset, reward, and 2D gaze position. **D.** Left: Histogram of the percent variance explained by the best model (open black) and the single best task variable (solid gray, quantified as the reduction in variance explained after removing that variable from the model). Right: Similar to left, computed using the best variable among other variables. **E.** Left: Histogram of the fraction of neurons tuned to different categories of task variables across all 32 recording session (including the 3 sessions included in the main results). A neuron was considered as tuned to a category if it was tuned to at least one variable from that category (e.g. tuned to sensory if it is tuned to either linear or angular velocity). Hand videos were not processed for recording sessions other than those presented in the main results. Instead, event-related temporal kernels were fit using the time of onset-of-movement and end-of-movement in lieu of the two leading PCs of hand speed. The rationale is that the hand speed components were large almost exclusively around those two time points and we verified separately that temporal kernels fit to hand speed were similar to kernels fit to onset and end of movement. Right: Similar to left, for other variables. **F.** Temporal filters associated with target onset for the set of neurons for which the filter peak fell within the period when the target was ON (within yellow window, top) and OFF (outside yellow window, bottom) respectively. Red line denotes a gain of 1 (no tuning). **G.** Tuning functions of neurons significantly tuned to horizontal (top) and vertical (bottom) eye positions, plotted as 'gain fields'. Red line denotes a gain of 1 (no tuning).

---

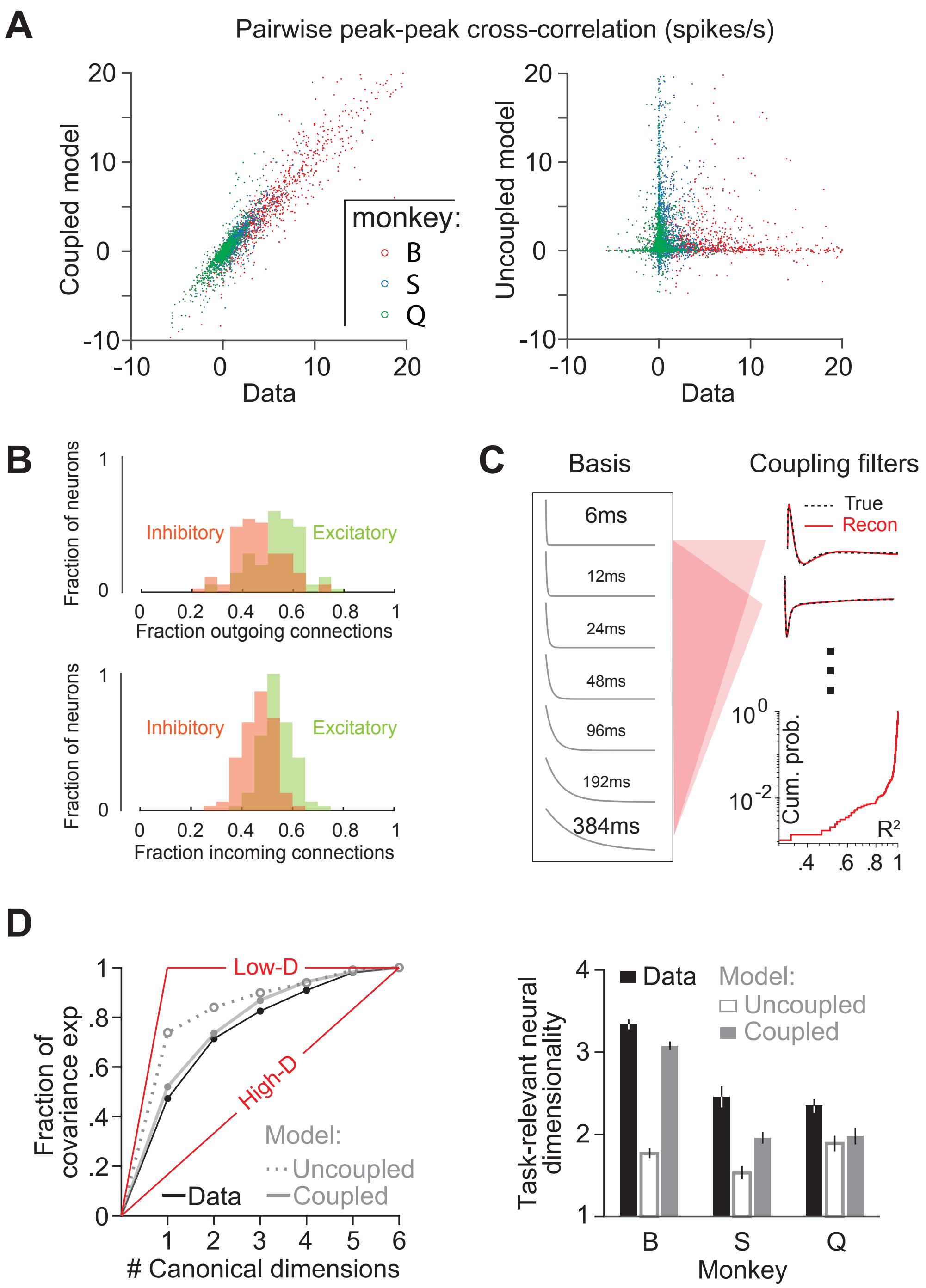
A
Pairwise peak-peak cross-correlation (spikes/s)
Coupled model
Data
monkey:
B
S
Q
Uncoupled model
Data
B
Fraction of neurons
Inhibitory
Excitatory
Fraction outgoing connections
Fraction incoming connections
C
Basis
Coupling filters
6ms
12ms
24ms
48ms
96ms
192ms
384ms
True
Recon
Cum. prob.
R²
D
Low-D
High-D
Fraction of covariance exp
### Canonical dimensions
Model:
Uncoupled
Coupled
Data
Task-relevant neural dimensionality
Monkey

*Figure S4:* **A.** Left: Comparison of the peak-peak cross-correlation value between pairs of neurons in data (abscissa) and the model with coupling between neurons (ordinate). Right: Similar to the left panel, but shown for the model without coupling. Red, Blue, and Green datapoints denote neuronal pairs from monkeys B, S, and Q respectively. **B.** For the purpose of classification, coupling filters with mean gain greater than unity were taken to be excitatory, and those less than unity were taken to be inhibitory. Top: Histogram of the fraction of outgoing connections that were excitatory (green) and inhibitory (red) across the population of neurons. Bottom: Similar to the top panel, but showing the fraction of incoming excitatory/inhibitory connections. **C.** The set of exponential basis functions of the form exp$(-t/\tau)$ used to approximate the best-fit coupling filters by linear regression. The numbers denote the time constants ($\tau$). Two example coupling filters (black dashed) and their reconstructions (red) are shown on the right. Exponential basis functions provided excellent reconstructions for all coupling filters (95% confidence interval $R^2$ = [.92 .99]) **D.** Left: Cumulative fraction of covariance between canonical task variables and canonical neural response for data (black) and simulated model response (gray), as a function of the number of canonical dimensions. Canonical correlations were computed by considering six task variables (see Figure 3D). Red lines denote extreme possibilities where covariance is either entirely concentrated within one dimension or distributed uniformly across six dimensions. Right: Task-relevant neural dimensionality computed from the covariance spectrum (Methods) computed from data and model neural responses for individual monkeys.

---

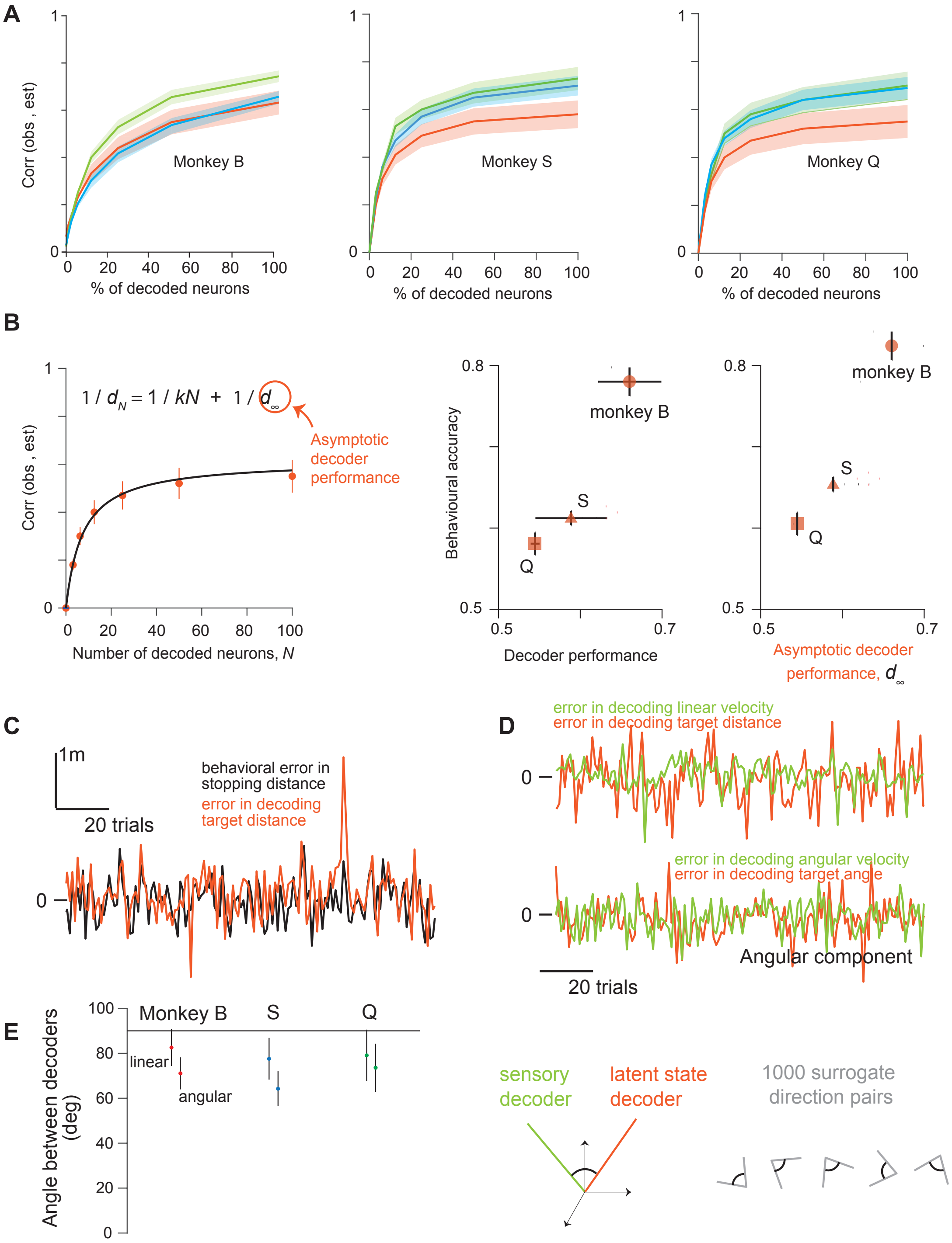

A
Corr (obs , est)
Monkey B
Monkey S
Monkey Q
% of decoded neurons
B
$1 / d_N = 1 / kN + 1 / d_\infty$
Asymptotic decoder performance
Number of decoded neurons, N
Behavioural accuracy
monkey B
S
Q
Decoder performance
Asymptotic decoder performance, $d_\infty$
C
1m
20 trials
behavioral error in stopping distance
error in decoding target distance
D
error in decoding linear velocity
error in decoding target distance
error in decoding angular velocity
error in decoding target angle
Angular component
20 trials
E
Angle between decoders (deg)
Monkey B
S
Q
linear
angular
sensory decoder
latent state decoder
1000 surrogate direction pairs

*Figure S5:* **A.** Decoder performance as a function of the percentage of neurons sampled from the population for monkey B (left), S (middle), and Q (right). Linear and angular components were qualitatively similar and have been averaged for ease of visualization. **B.** Left: Asymptotic performance of the latent state decoder determined by parametric extrapolation. Middle and right: The relationship between behavioral accuracy and the raw decoding performance (middle, reproduced from main text Fig. 5C) or the asymptotic performance (right). **C.** Trial-by-trial fluctuations in behavior closely followed the latent state decoder performance. The timescale of these fluctuations is rapid suggesting that the correlation could not be explained by slowly varying factors such as level of arousal or wakefulness. **D.** Similar to panel C, but showing fluctuations in sensory (green) and latent state decoders (orange) for an example monkey. Top and bottom panels show results in linear and angular components respectively. **E.** Left: The angle between sensory and latent state decoding directions for both linear and angular components of all three monkeys. Errorbars denote 95% confidence intervals estimated by bootstrapping. Horizontal line corresponds to orthogonality (90 degrees). Right: For each component of each monkey, surrogate direction pairs with the same angles as data were used to generate null distribution of correlation coefficients (Fig. 5E – right).

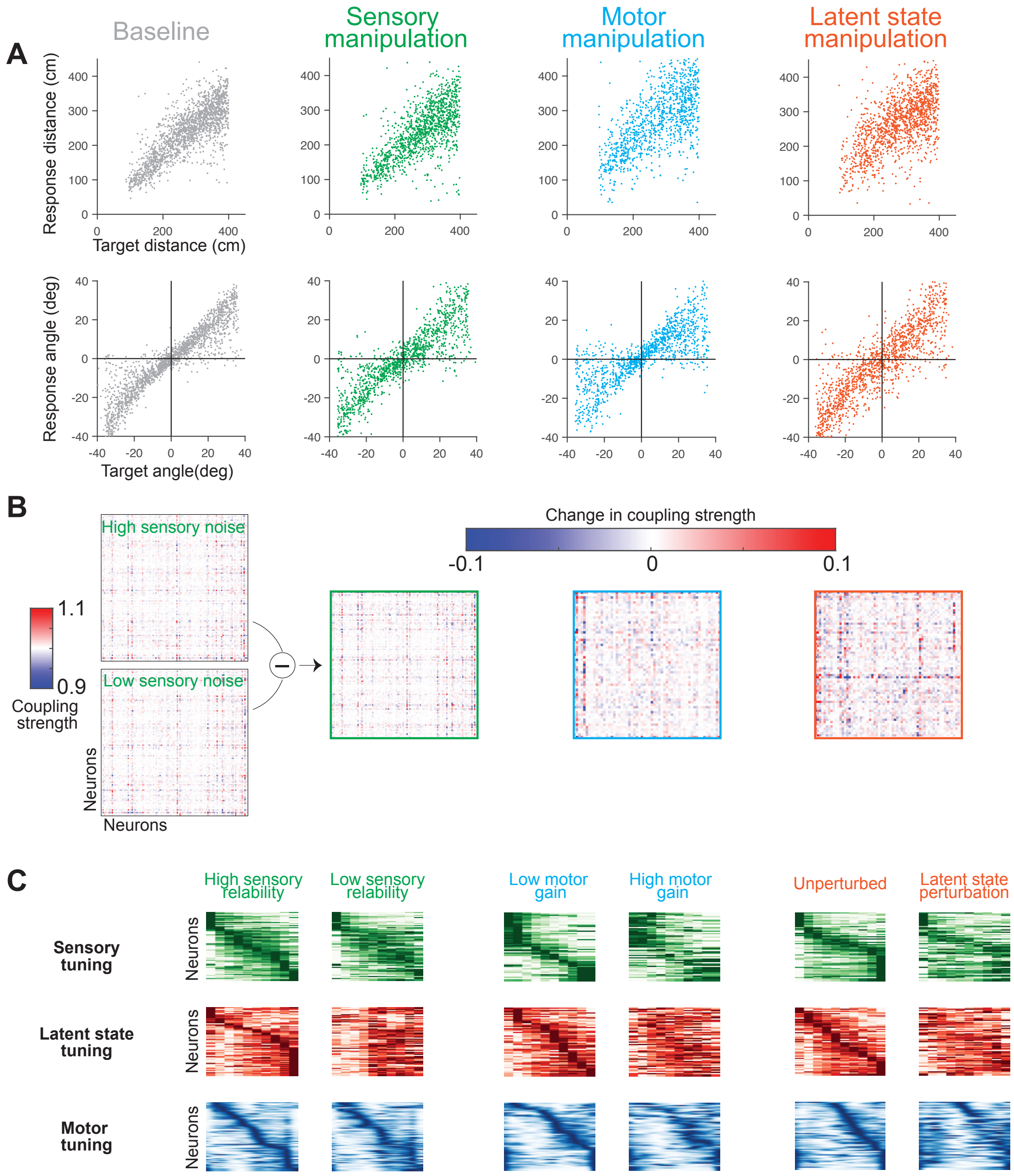
A
Baseline
Sensory manipulation
Motor manipulation
Latent state manipulation
Response distance (cm)
Target distance (cm)
Response angle (deg)
Target angle(deg)
B
High sensory noise
Low sensory noise
1.1
0.9
Coupling strength
Neurons
Neurons
Change in coupling strength
-0.1
0
0.1
C
High sensory reliability
Low sensory reliability
Low motor gain
High motor gain
Unperturbed
Latent state perturbation
Sensory tuning
Latent state tuning
Motor tuning
Neurons

*Figure S6:* **A.** Top: Comparison of the radial distance of the response against radial distance of the target across a subset of trials from various conditions (grey – baseline, green – manipulate sensory reliability, cyan – manipulate motor gain, red – manipulate latent state dynamics). Bottom: Angular eccentricity of the response versus target angle for the same four conditions. Data from all three monkeys are included in these plots. **B.** Matrix of coupling strengths (determined by the area under the coupling filter) had roughly the same structure in both baseline and manipulated trials, but were not identical. The difference between coupling matrices fit to data from baseline and manipulated trials is shown for one example session of each of the three manipulations. **C.** Effect of task manipulation on neural tuning to sensory (top row), latent state (middle row), and motor (bottom row) variables. In each panel, the horizontal and vertical axes denote stimulus value and neurons respectively, and color denotes normalized firing rate (dark is high). For each manipulation, neurons are sorted based on the preferred stimulus feature in the baseline condition (left column) and this same ordering is used to plot the tuning functions during task manipulation (right column). To keep the visualization simple, only one of the two dimensions is shown for each variable (linear velocity for sensory, target distance for latent, and the first PC of hand motion for motor. Results were qualitatively similar for the angular dimension. Stimulus range (values omitted for the sake of clarity) is same as Figure 3E for all variables except motor where a smaller time range around zero is shown to facilitate comparison of the response timing.

---

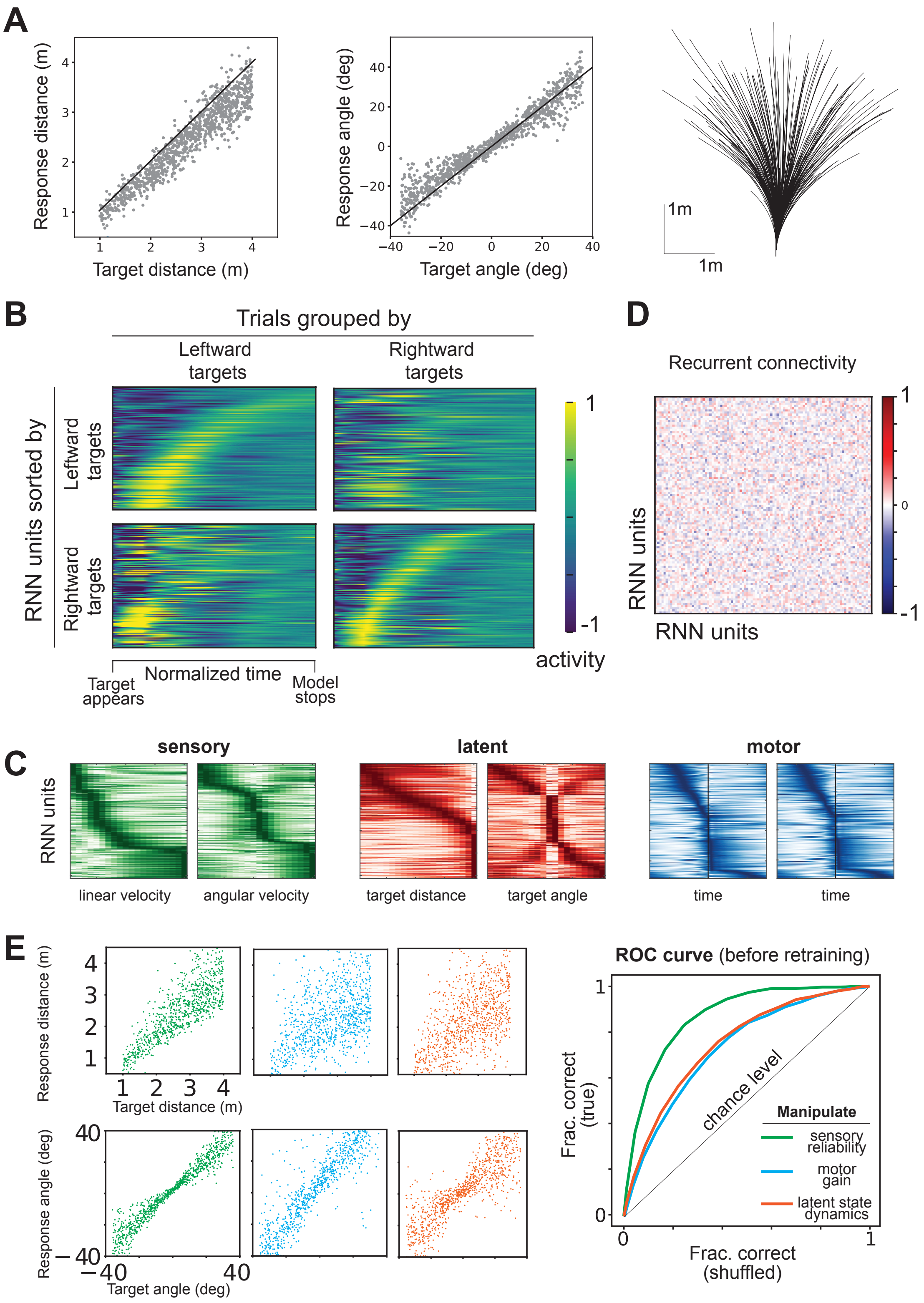
A
Response distance (m)
Target distance (m)
Response angle (deg)
Target angle (deg)
1m
1m
B
Trials grouped by
Leftward targets
Rightward targets
RNN units sorted by
Leftward targets
Rightward targets
Normalized time
Target appears
Model stops
activity
D
Recurrent connectivity
RNN units
RNN units
C
sensory
latent
motor
RNN units
linear velocity
angular velocity
target distance
target angle
time
time
E
Response distance (m)
Target distance (m)
Response angle (deg)
Target angle (deg)
ROC curve (before retraining)
Frac. correct (true)
Frac. correct (shuffled)
chance level
Manipulate
sensory reliability
motor gain
latent state dynamics

*Figure S7:* **A.** Left: Comparison of the radial distance of the response against radial distance of the target across a subset of trials from the trained recurrent neural network. Middle: Angular eccentricity of the response versus target angle. Black dashed lines have unity slope. The starting position was taken as the origin. Right: Spatial trajectories on a subset of trials. **B.** Peak-normalized response of RNN units calculated by averaging across the set of trials with leftward (left panels) or rightward (right panels) targets. Units are sorted according to the timing of their peak response observed in the set of trials with leftward (top panels) or rightward (bottom panels) targets. Time was rescaled based on the trial duration before trial-averaging and the resulting response profile of each unit was subsequently normalized by the peak activity observed in the condition used for sorting. This RNN was trained with a regularizer that penalized high activity and encouraged smooth dynamics. **C.** Peak-normalized tuning functions of RNN units, sorted according to the peak feature. Labels and other conventions are similar to Figure 3F. **D.** Recurrent connectivity matrix of the trained recurrent neural network showing roughly equal number of excitatory and inhibitory connections. **E.** Left: Top panels show a comparison of the radial distance of response against the target for the three manipulations tested on the RNN before retraining. Bottom panels show a comparison of the angular eccentricities. Right: ROC curves comparing the generalization performance for the three manipulations. The network showed some generalization to all manipulations, but was most robust to adding sensory noise.

---
